# Historical recombination variability contributes to deciphering the genetic basis of phenotypic traits

**DOI:** 10.1101/792747

**Authors:** Carlos Ruiz-Arenas, Alejandro Cáceres, Marcos López, Dolors Pelegrí-Sisó, Josefa González, Juan R. González

**Affiliations:** Instituto de Salud Global de Barcelona, Doctor Aiguader 88, Barcelona 08003, Spain; Universitat Pompeu Fabra (UPF), Barcelona, Spain; CIBER Epidemiología y Salud Pública (CIBERESP), Barcelona, Spain; Genetics Unit, Universitat Pompeu Fabra, Barcelona 08003, Spain; Centro de Investigación Biomédica en Red de Enfermedades Raras (CIBERER), Barcelona 08003; Institute of Evolutionary Biology, Passeig Marítim de la Barceloneta 37-49, Barcelona 08003, Spain

## Abstract

Recombination is a main source of genetic variability. However, the potential role of the variation generated by recombination in phenotypic traits, including diseases, remains unexplored as there is currently no method to infer chromosomal subpopulations based on recombination patterns differences. We developed *recombClust*, a method that uses SNP-phased data to detect differences in historic recombination in a chromosome population. We validated our method by performing simulations and by using real data to accurately predict the alleles of well known recombination modifiers, including common inversions in *Drosophila melanogaster* and human, and the chromosomes under selective pressure at the lactase locus in humans. We then applied *recombClust* to the complex human 1q21.1 region, where nonallelic homologous recombination produces deleterious phenotypes. We discovered and validated the presence of two different recombination histories in these regions that significantly associated with the differential expression of *ANKRD35* in whole blood and that were in high linkage with variants previously associated with hypertension. By detecting differences in historic recombination, our method opens a way to assess the influence of recombination variation in phenotypic traits.

## Introduction

Recombination plays a central role in adaptation and evolution, and its influence in human disease is becoming increasingly clear [1]. During the last decade, our understanding of genome-wide recombination rates and landscape has been greatly increased by the resolution and power of high-throughput data and analysis methods on population samples. Methods that extract recombination signals from linkage between SNPs have been instrumental [2–6]. However, despite these great advances, the outstanding question on how recombination variability influences phenotypes has lagged behind as there has not been a method to measure recombination variation between individuals for large association studies. A large body of theoretical work, initiated by Nei [7], has explored the conditions under which the variability of general recombination modifiers evolve [8, 9] yet empirical studies that link recombination variability in a genomic region with phenotypic traits and fitness are restricted to already known specific modifiers, such as inversions or specific polymorphisms [10–12]. In this context, we developed *recombClust*, a pioneer method to detect recombination variability between chromosomes by inferring the differences in recombination histories within a genomic region.

Recombination produces offspring chromosomes with new combinations of maternal and paternal DNA material at each side of a recombination event [13]; making it a main source of novel genetic diversity. At the population level, when multiple recombination events have occured between two genomic markers, the linkage between them decreases and a random association is then observed. Historic recombination patterns were thus successfully extracted from the linkage between dense SNP markers, strongly matching direct observations on recombination events in sperm samples [14]. Because linkage methods are population-based estimates, they have been intensely used to compute accurate recombination rates and landscapes in large population samples but, at the same time, have also been disregarded in their ability to detect recombination variation between individuals [15], *i.e*. used to infer groups of chromosomes with different recombination histories in a genomic region. However, latent variable mixture models can be incorporated to linkage methods to detect the underlying mixture of chromosome subpopulations, characterized by different recombination patterns. We, therefore, hypothesized that in a genomic region where the recombination frequency and location are modified in a subpopulation of chromosomes, the chromosomes can be grouped according to consistent recombination histories within the region. The detected chromosome groups could then be tested for association with phenotypes, allowing the use of large cohorts to study the phenotypic effects of recombination variability in the genomic region.

Here, we proposed a method that leverages chromosomal differences in linkage patterns in a genomic region to classify the chromosomes of a population into groups with different recombination histories. The method, named *recombClust*, comprises two steps. First, it fits a mixture model for each pair of SNP blocks within a genomic region to classify chromosomes into those with a history of high recombination or high linkage between the blocks; second, it tests the consistency of the chromosomes’ classification across all the mixture models *i.e*. all SNP block pairs. Chromosome groups with different recombination histories are thus called by the chromosome’s classification into the consistent recombining groups. By estimating the proportion of chromosomes with historic recombination at a given point in the region, the recombination pattern for each chromosome subpopulation can be reconstructed.

We tested the performance and adequacy of the method using numerous simulated scenarios and demonstrated its ability to detect known recombination modifiers with their correct recombination patterns using real data for *Drosophila melanogaster* and humans. Finally, we used the method to i) detect and validate chromosome subpopulations with different historic recombination at 1q21.1, a genomic region at risk of deleterious rearrangements leading to the thrombocytopenia-absent radius (TAR) syndrome [16, 17], and ii) to associate the chromosome groups with changes in gene expression in blood. The method was implemented in a computationally efficient tool, compatible with Bioconductor’s packages and the variant call format (VCF). The development version is available at https://github.com/isglobal-brge/recombClust and the final version will be available in Bioconductor.

## Results

We implemented *recombClust*, a method to classify chromosomes into groups with different recombination histories across a genomic region (Figure 1). The method comprises two steps. First, for each pair of SNP blocks in a genomic region, it fits a mixture model of two chromosome groups (*recomb/linkage*), one in which chromosomes display random association between the blocks (*recomb*) and the other where the blocks are found in complete association (*linkage*). Second, *recombClust* classifies chromosomes into subpopulations (A/B) based on a consensus clustering across all the mixture models fitted along the genomic region. The chromosome groups A/B are the subpopulations associated with different recombination histories, which can be reconstructed from the proportion of chromosomes in the *recomb* group at each point across the genomic region. The underlying chromosome substructructure (A/B) can be used in downstream analysis, such as transcription and methylation profiling or association with phenotypes.

**Figure 1:**
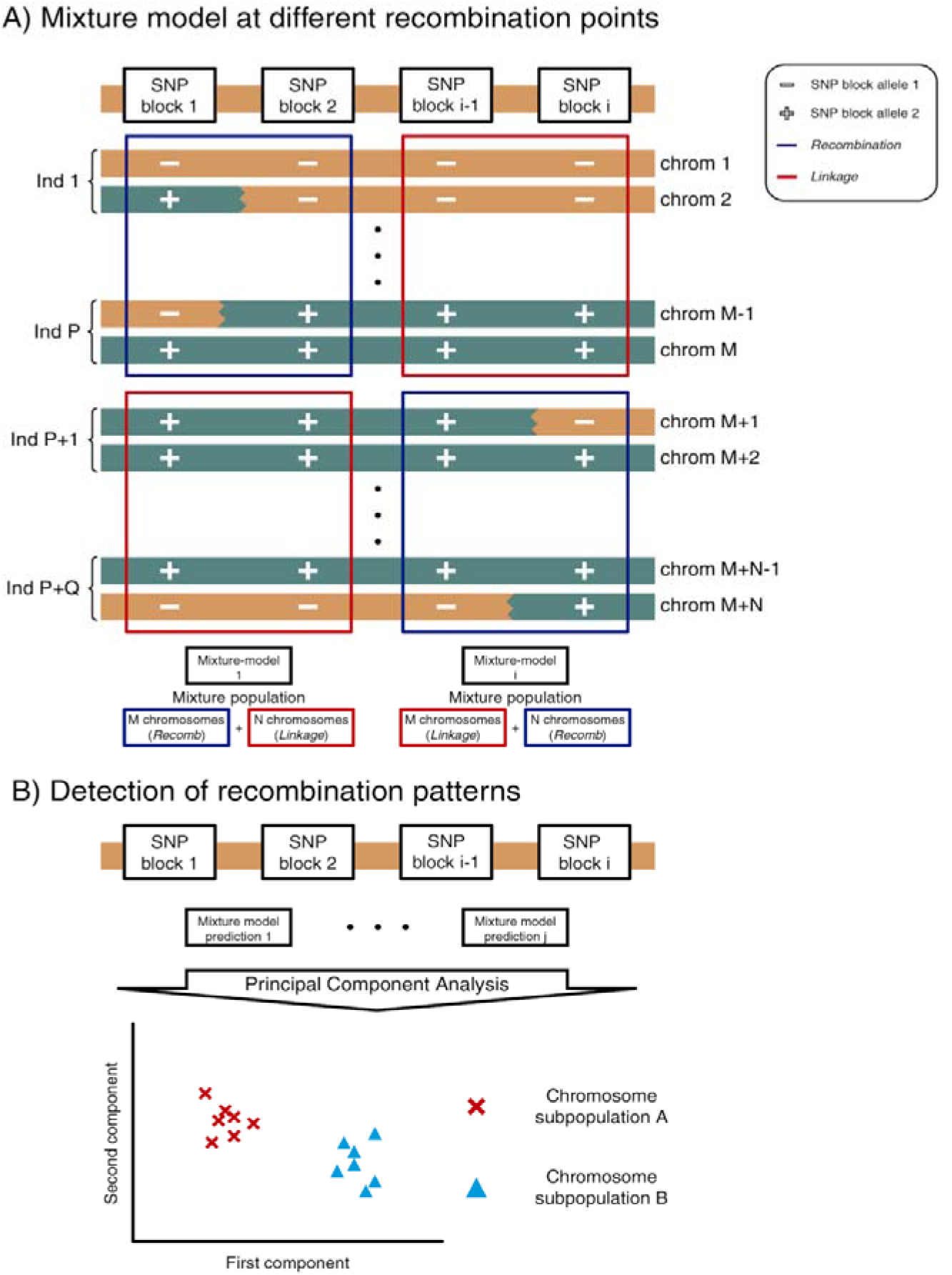
*recombClust* scheme. *recombClust* is a method to classify chromosomes into underlying recombination patterns using SNP data, it comprised two steps illustrated at the top and bottom of the figure. A) Mixture model fitting at different recombination points. Colours represent two main haplotypes. In the chromosome subpopulation on the left, there is recombination between the blocks in P chromosomes as they have all possible allele combinations between the SNP blocks 1/2, illustrated in geometrical figures. Q chromosomes are in high linkage. Whereas, for the population on the right, the opposite situation is observed for SNP blocks *i-1/i*. At each point the population is a mixture of P and Q chromosomes into the *recomb* and *linkage* groups. B) Chromosome classification into recombination patterns A and B. The mixture models provide a classification at each point, the first PCs of the classification matrix across all mixture models along the genomic region can detect clusters of chromosomes for which their classifications are similar, and therefore share similar recombination patterns. Each chromosome is assigned to a recombination subpopulation. The recombination pattern for each chromosome subpopulation can be reconstructed from the proportion of chromosomes in the *recomb* group at each point in the genomic region (not shown).

### Modeling the mixture of chromosomes under recombination and linkage

We developed a mixture model to split the chromosomes of a population into those showing high recombination and those showing high linkage history between two SNP blocks (Methods). Figure 1A illustrates two instances where the mixture model is fitted at two different points in a genomic region. For illustration purposes, only two alleles are shown at each SNP block (*+*,-). The first recombination point is tested by blocks *1/2* where *M* chromosomes are in the *recomb* group showing random association between the blocks and *N* chromosomes are in the *linkage* group showing maximum linkage between the blocks. The other point is tested by blocks *i–1/i* where now the *N* previous chromosomes belong to the *recomb* group and the previous *M* chromosomes to the *linkage* group. Note that although the model at one point can be ambiguous for some chromosomes (i.e. *chrom 1* at *1/2* in Figure 1A), the final chromosome classification into subpopulations consistent with specific recombination patterns is robust when considering other points in the region (*chrom 1* at *i–1/i*), as explained in the following section.

We simulated multiple datasets representing a SNP block pair that flanked one recombination point for a group of chromosomes (*recomb*) but remained in linkage for a second group (*linkage*), see Methods section. We first evaluated how the proportion between *recomb* and *linkage* populations affected the accuracy of the model to correctly classify the chromosomes, varying the proportion between 0.1 and 0.9. We observed that the mixture model had high accuracy (>80%) across all the proportion range, being optimal, as expected, when the mixture was small, *i.e*. the mixture frequency approached to 1 or 0 (Sup Figure 1A). We also observed that the model was robust under different initializations of the mixture frequency (Sup Figure 1B). Overall, our simulations showed that the mixture model was able to robustly split the chromosomes into two groups, one with null LD (*recomb*) and other with full LD (*linkage*) between a pair of 2-SNP blocks.

We then evaluated the accuracy of the model under different within and between SNP block variabilities, using a fix scenario with a 0.5 proportion of mixture between the *recomb* and *linkage* groups. To test SNP block variability, we simulated multiple two-SNP block pairs, flanking a recombination point, and determined the haplotypes across the blocks. We varied the number of SNP alleles that were different between the most frequent *recomb* and *linkage* haplotypes. We thus assessed the extent to which the accuracy of the model was affected by increasing differences in haplotype variability between the groups. We observed that the mixture model had an accuracy of 75% when most frequent haplotypes were shared between groups, and topped to 90% when the difference between the haplotypes was given by only one SNP allele (Sup Figure 1C). This suggests a substantial accuracy when the differences in mutation frequency between the groups are small, which, in addition, can be boosted by the presence of one SNP allele that associates with one of the groups. We then assessed the influence of within block variability on model accuracy. For an scenario of full linkage of the SNPs within all blocks, which reduces to having blocks of 1 SNP, the accuracy dropped to ^~^60%, showing that larger and more variable SNP blocks increase model’s accuracy (Sup Figure 1D).

### Classifying chromosomes into different recombination histories within a genomic region

The second step of *recombClust* is a consensus clustering of mixture models at numerous points along a genomic region to classify chromosomes into consistent recombining groups (A/B) (Figure 1B). Within the region, all possible SNP blocks pairs are tested such that they do not overlap and are at a maximum distance of 10kb. For each block pair, a mixture model is fitted and the chromosomes classified into the *recomb* and *linkage* groups. Because at one point in the region, chromosomes in *recomb* can be in *linkage* at another point, a consistent classification over the mixture model predictions was considered. For this step, we applied a clustering method (k-means) on the first PCA components of the model prediction variables obtained from the mixture models fitted along the region. The clusters identified were then considered as chromosomes with similar recombination patterns within the region. Mixture model classification across the region was used to reconstruct the pattern of classification proportion into the *recomb* and *linkage* groups, this pattern was then compared with the recombination patterns obtained by other linkage based methods, which are applicable only when the chromosome subpopulation A/B are initially known.

We used simulations to test whether the number of chromosomes and the number of recombination points affected the accuracy of *recombClust* to identify subpopulations of chromosomes with different recombination patterns. We thus simulated datasets representing SNP block pairs that flanked multiple recombination points. We simulated two kinds of populations: (1) a mixture population, where one subpopulation (A) belonged to the *recomb* group in half of the points and to the *linkage* group in the other half while a second subpopulation (B) belonged to the *linkage* and *recomb* groups, respectively; and (2) a single population where all chromosomes belonged to the same recombination groups across all recombination points.

First, to assess false discovery rate and statistical power, we selected several scenarios changing the number of chromosomes per population (from 20 to 60) and the number of recombination points (from 10 to 100). In all cases, we performed a PCA to the classification matrix given by the mixture model probabilities of belonging to a *recomb* group at each SNP block pair (Figure 1B). Then, using k-means, we clustered the first two PC components in two groups and considered that *recombClust* detected differences in recombination patterns when the average silhouette value of the clustering was higher than 0.7 [18] (Sup Figure 2). We observed that under single population simulations, *recombClust* had a false discovery rate < 0.05 for recombination points > 70 and for all the number of chromosomes considered (>20). In addition, the power to detect different recombination patterns for simulations of chromosomes with two different recombination histories achieved 80% for > 25 chromosomes and for differences in historical recombination in > 16 points (Sup Figure 3).

Second, to confirm that the model detected differences in recombination histories rather than allele differences, we compared *recombClust* classification with that of a PCA on the simulated genotypes. For a simulation with chromosome mixture, we observed a neat separation of the chromosome subpopulations (Sup Figure 4A) with *recombClust*, which we did not observe for allele differences.

### *recombClust* accurately classifies inversion status based on differences in historic recombination

The alleles of polymorphic inversions differ in the recombination histories inside the inverted region because recombination is suppressed in heterokaryotypes [3]. We, therefore, asked the extent to which the inversion alleles, being strong recombination modifiers, could be inferred by recombination differences using *recombClust*. We evaluated the method’s performance to predict simulated inversions from the coalescent simulator *invertFREGENE* and tested its accuracy to classify common inversions in *Drosophila melanogaster* and humans. Using *invertFREGENE* [19], we simulated inversions with different lengths (from 50Kb to 1Mb) and frequencies (from 0.1 to 0.9) and tested the prediction accuracy of chromosome classification into their inversion alleles. We observed accuracy greater than 90% for inversions larger than 250Kb (Sup Figure 4B). As expected, accuracy for short inversions was lower as they presented fewer recombination points. *recombClust*’s mean accuracy was higher (95%) for inversion frequencies between 0.2 and 0.8 (Sup Figure 5) but did not correlate with the inversion’s age (r = 0.02, p-value = 0.19) (Sup Figure 6).

We then used *recombClust* to determine whether the alleles of three common polymorphic chromosomal inversions in *Drosophila melanogaster* (In(2L)t, In(2R)NS and In(3R)Mo) could be determined from different recombination histories. We ran *recombClust* on genome-wide SNP data of 205 lines derived from Raleigh, USA population, comprised in the Drosophila Genetics Reference Panel (DGRP2)[20, 21] and compared the inferred recombining subpopulations with the experimental inversion alleles of the lines. For all the inversions, we observed clear clustering in the first PC component of the mixture classification matrix (Figure 2) that resulted in a 98% match with the inversion alleles, when a k-means clustering was applied. Likewise, we compared the *recombClust* calling of human inversions at 8p23.1 and 17q21.31 with the experimental inversion genotypes, as obtained from the *invFEST* repository [22] for the European subjects of the 1000 Genomes Project. Using SNP-phased data, we found that *recombClust* neatly separated inverted and standard chromosomes (Figure 2) in the first PC component of the mixture classification matrix. The k-means clustering of the first PC accurately matched the experimental inversion-alleles (8p23.1: 100%, 17q21.31: 99.3%). Overall, these results showed that recombination substructure can reliably identify the inversion alleles of some common inversions in two different species.

**Figure 2:**
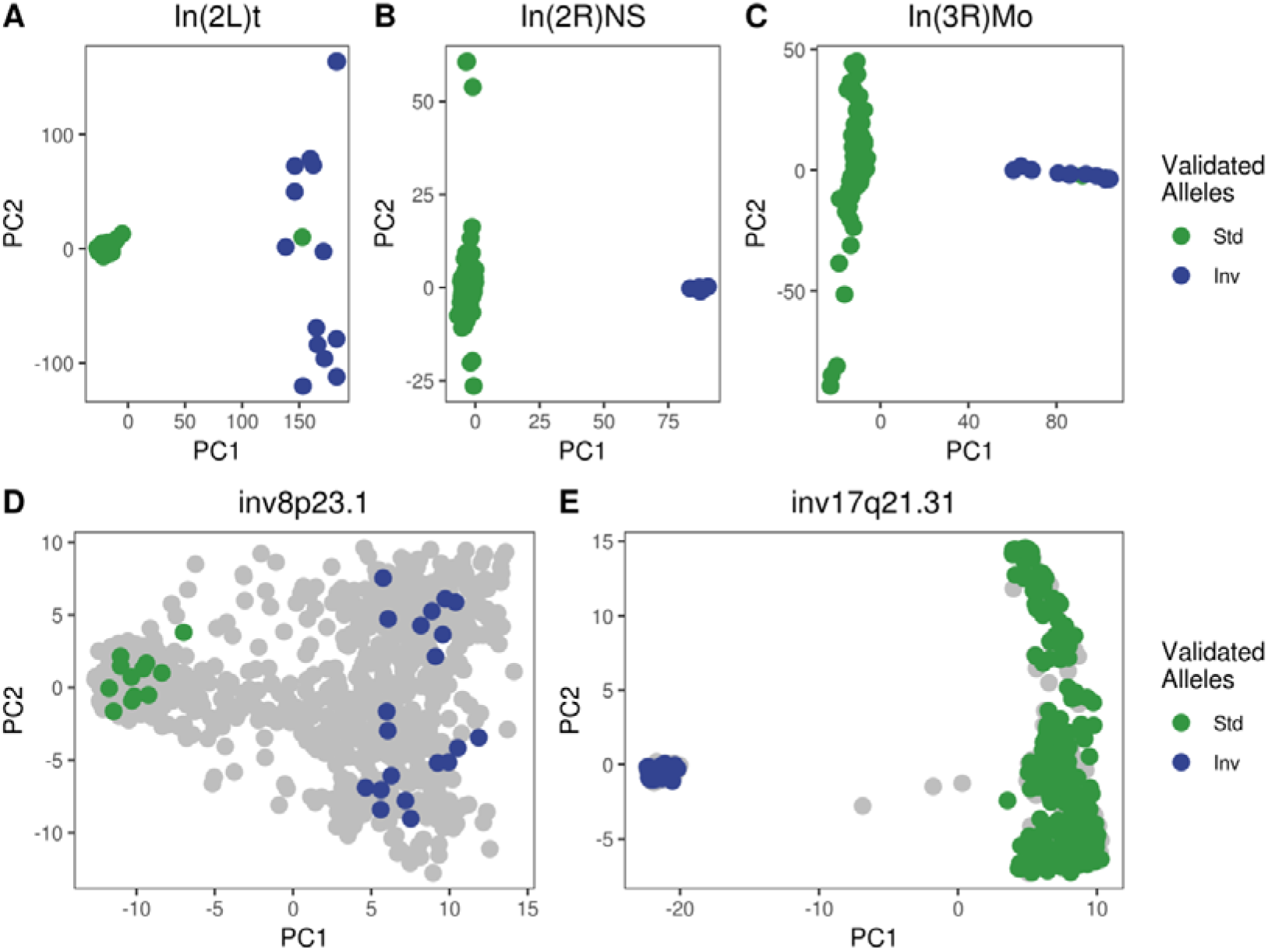
PCAs of *recombClust* probabilities for chromosomal inversions in *Drosophila Melanogaster* and human. First two principal components of chromosomes, derived from the recombination classification at multiple recombination points along different inverted regions. Each point is a chromosome. Clusters mapping the inversion status in both *Drosophila* and human inversions are clearly observed. Chromosomes with known inversion genotypes are coloured (green: standard, blue: inverted). A-C) *Drosophila* inversions in DGRP2 lines. D-E) Human inversions in the European individuals of the 1000 Genomes project.

To test whether *recombClust* classification reflects differences in historical recombination rates, we compared the recombination pattern obtained along the longest human polymorphic inversion 8p23.1 with *recombClust* with the recombination rates estimated with FastEPRR [23] independently obtained for each inversion allele (Figure 3). Remarkably, we observed that the inferred proportion of chromosomes in the *recomb* population across the genomic region accurately captured the underlying recombination patterns obtained by *FastEPRR* for each of the 8p23.1 inversion alleles. We also observed that the largest differences in recombination proportion were obtained near the recombination hotspots obtained by Alves et al [3] (Figure 3).These results confirmed that the chromosome subpopulations identified by *recombClust* are clearly mapped to different recombination histories.

**Figure 3:**
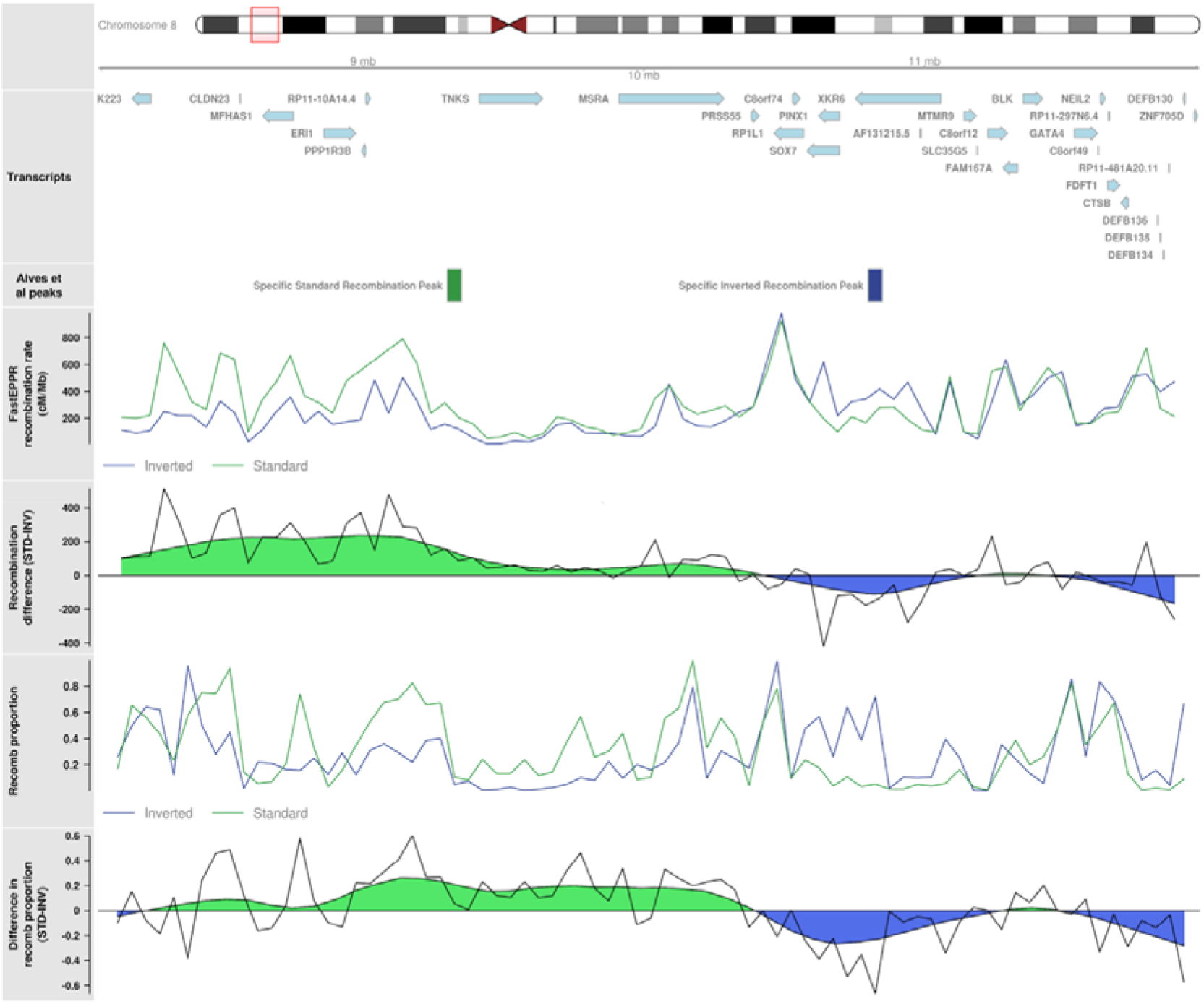
Underlying recombination patterns in human inversions 8p23.1 for the European individuals of the 1000 Genomes Project. Ideogram for the 8p23.1 inverted region showing the transcripts in the region. *Alves et al* peaks: approximate location of recombination hotspots for standard or inverted chromosomes identified by Alves and colleagues [3]. *FastEPRR recombination rate:* recombination rate obtained from *FastEPRR* independently for standard and inverted chromosomes. *Recombination difference:* raw and smoothed difference in recombination rates between standard and inverted chromosomes as computed from *FastEPRR. Recomb proportion:* proportion of chromosomes belonging to *recomb* population in the chromosome subpopulations detected by recombClust, which accurately predicted inversion status. *Difference in recomb proportion:* raw and smoothed difference in the proportion of chromosomes belonging to *recomb* population in inverted and standard chromosomes, as predicted by *recombClust*.

### *recombClust* detects recombination histories associated to ancestral differences

Modifiers of historical recombination patterns include numerous processes other than inversions that can act simultaneously on the same genomic region. In particular, differences in historical recombination patterns between ancestries can derive from random differences in the occurrence of recombination events or from the emergence of hotspot differences regulated by ancestry-specific alleles [24]. As such, we asked the extent to which differences between human populations could also be detected in loci already under the influence of inversion alleles. We, therefore, used *recombClust* to detect the modifier alleles in the loci corresponding to the human inversions at 8p23.1 and 17q21.31 for all the individuals in the 1000 Genomes Project, covering four different continental populations [25]. We inspected the first two PC components of the mixture model predictions for inv-8p23.1 (Sup Figure 7), and observed multiple clusters, in which chromosomes segregated both by inversion status and ancestry. However, for inv-17q21.31, the additional clusters observed in the standard allele did not map to ancestral differences. The observations on both human inversions confirmed that that clusters identified in the first PCs of the mixture model predictions can be interpreted as non-recombining chromosome groups that differ in inversion status, ancestry, or other unobserved factors that suppress recombination between the groups, such as copy number variants likely segregating in standard chromosomes at 17q21.31 [26].

### *recombClust* detects recombination histories associated with selection

Chromosomes with advantageous alleles show a decrease in recombination around the locus under selection. While selection, like demography, does not have a direct influence on the biological process of recombination, they modulate the historical recombination patterns [5]. Therefore, we asked whether *recombClust* was able to detect chromosomes under selection and recover their recombination patterns. We studied the *LCT* locus, a human locus known to be under positive selection for lactase tolerance, as defined in PopHumanScan (chr2:135770000-136900000, hg19) [27]. We aimed to detect the underlying chromosomes under selection and their recombination pattern in the *LCT* locus, for the European individuals of the 1000 Genomes Project. We observed two chromosomes subpopulations (A/B) by clustering the first PC components of the mixture classification matrix (allele 1: 60.8%, allele 2: 39.2%) (Figure 4). Notably, chromosome allele 1 was the most frequent except for the Tuscany population (TSI) (Sup Table 1), the only European population which does not show marks of selection in the *LCT* locus, as reported in PopHumanScan [27]. We also observed a strong correlation between rs4988235 (C/T(-13910)), the SNP linked to lactose resistance, and the inferred subpopulation groups (r^2^ = 0.64), where the allele conferring lactose resistance (T) was very frequent in chromosome allele 1 (83%) and almost absent in chromosome allele 2 (<1%). The ability of *recombClust* to detect chromosomes under selection was further confirmed by the proportion of chromosomes in *recomb* along the locus for each chromosome subpopulation. As expected, we confirmed that recombination appeared flat in group A (under selection) but not in B (not under selection) across the *LTC* locus (Figure 4). We also recovered the recombination patterns independently obtained with *FastEPRR*, for each chromosome subpopulation. Recombination peaks for chromosome allele 2 were found between genes *R3HDM1* and *DARS* genes, matching previously reported recombination peaks [28].

**Figure 4:**
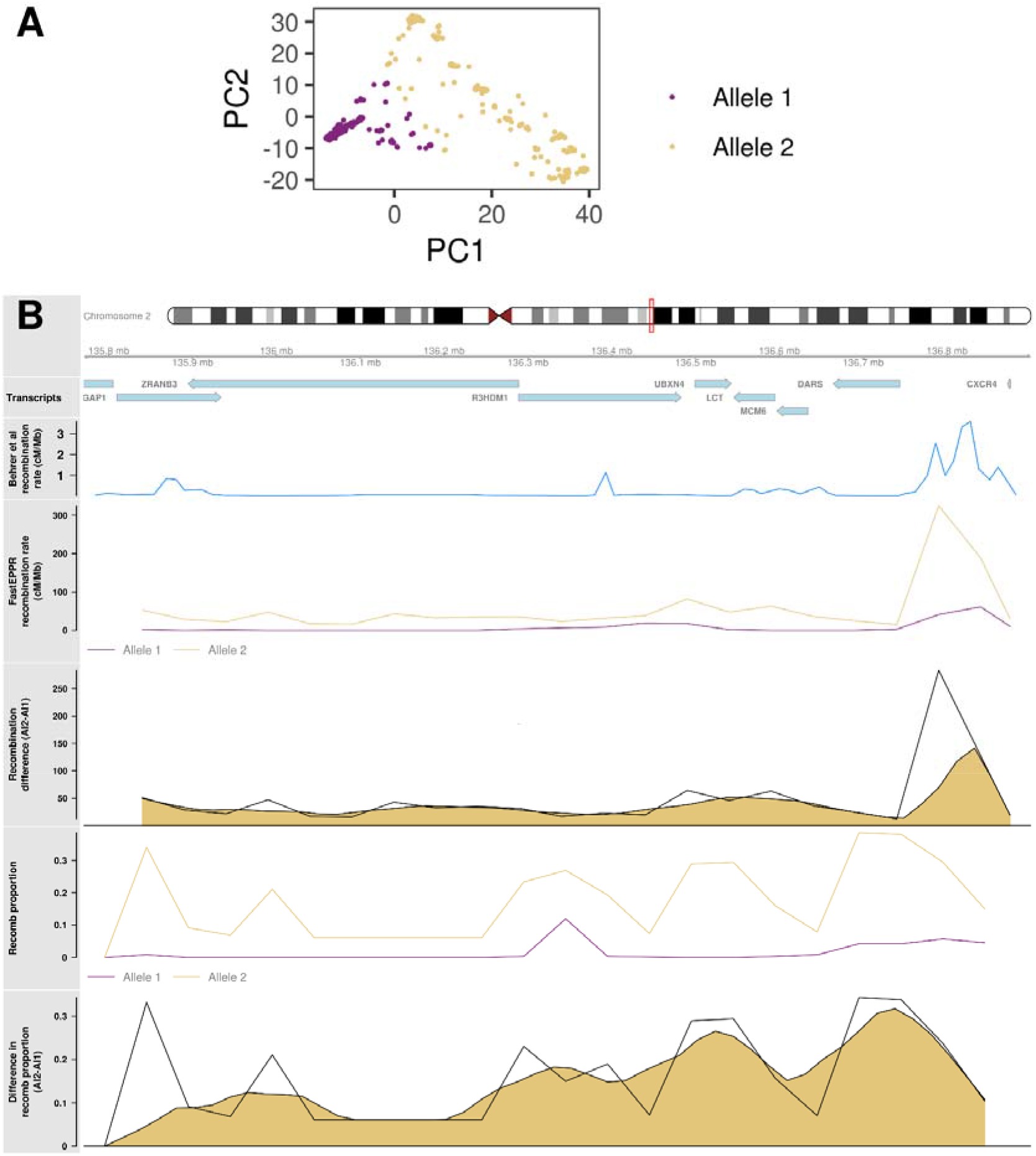
Underlying recombination patterns in the *LCT* locus. A) First two principal components of chromosomes, derived from the recombination classification at multiple recombination points along the *LCT* locus. B) Ideogram for the *LCT* locus under selection showing the genes in the region. *Bhérer et al recombination rate:* recombination rates reported by Bherer and colleagues [28]. *FastEPRR recombination rate:* recombination rate obtained from *FastEPRR* independently for chromosomes with alleles 1 and 2 detected by *recombClust. Recombination difference:* raw and smoothed difference in recombination rates between alleles 1 and 2 as computed from *FastEPRR. Recomb proportion:* proportion of chromosomes belonging to *recomb* population in the chromosome subpopulations with alleles 1 and 2, correctly predicting a flat pattern for the allele 1 that is under selection. *Difference in recomb proportion:* raw and smoothed difference in the proportion of chromosomes belonging to *recomb* population in in alleles 1 and 2.

### *recombClust* detects recombination differences in complex genomic regions

The region at 1q21.1 between chr1:145,399,075-145,594,214 (hg19) [29] is prone to various deleterious rearrangements by non-allelic homologous recombination (NAHR) at the numerous segmental duplications (SD) in the region [16]. The rearrangements include microdeletions leading to the thrombocytopenia-absent radius (TAR) syndrome and a range of multiple neurodevelopmental phenotypes caused by duplications and deletions distal to the TAR region [16]. As strong control of recombination is expected in regions regions at risk of NAHR during meiosis [30], we hypothesized that different recombination histories would be detectable in region and aimed to determine their functional correlates.

We ran *recombClust* across the region chr1:145.35-145.75Mb characterized by four blocks of segmental duplications. The most common deletion for the TAR syndrome is observed between the first and third block [31] while the smallest reported deletion was found between the second and third block [16] (Figure 5). We first analyzed the European individuals of the 1000 Genomes project and observed two clear clusters in the first two PCs of the classification matrix across mixture models. We defined two chromosome subpopulations (subpopulation 1: 80.9%, subpopulation 2: 19.1%) that were in Hardy-Weinberg equilibrium (*P* = 1) and thus confirmed our hypothesis for the presence of different recombination histories in the region. For each group, we estimated the recombination pattern given by the proportion of chromosomes in *recomb* (Figure 5), observing important differences between the groups. Notably chromosomes in subpopulation 2 had higher recombination proportion than those in subpopulation 1 along the region except for the small interval containing the genes *LIX1L* and *RBM8A*, the causative gene of TAR syndrome [29]. However, the highest differences in recombination proportions were observed between the third and fourth SD blocks, where subpopulation 1 showed null recombination; suggesting a stronger suppression of recombination for this group of chromosomes. We fully validated the chromosome subpopulations and their recombination patterns using the Whole Genome Sequencing data of 287 European individuals from the Genotype-Tissue Expression project (GTEx) (Figure 5). We thus obtained strong evidence for the existence of two recombination histories in the region.

**Figure 5:**
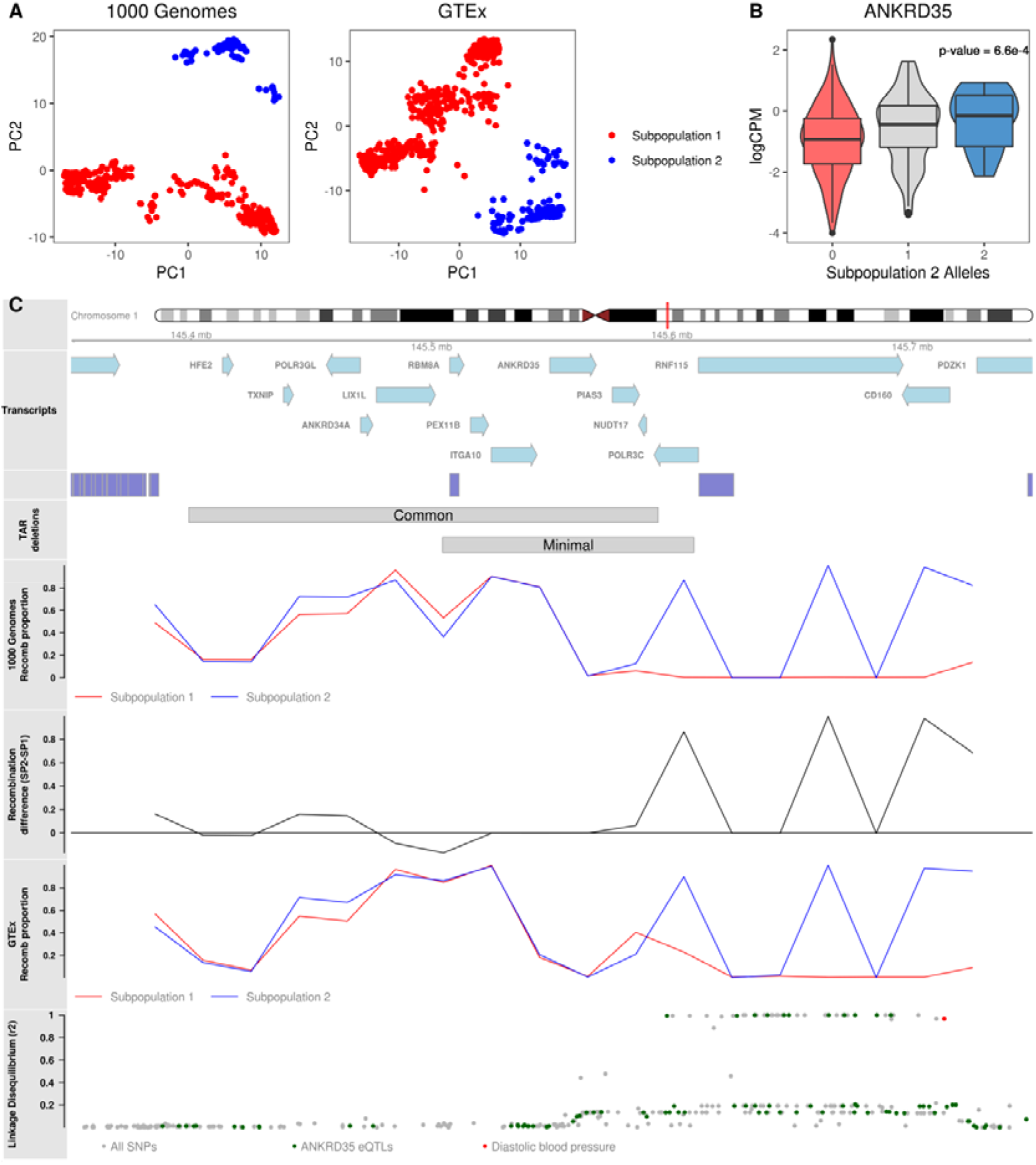
Underlying recombination patterns in the TAR syndrome locus. A) Chromosome subpopulations with different recombination pattern between the coordinates chr1:145.35-145.75 (hg19), as detected in the genomic data of the 1000 Genomes and GTEx projects. B) Transcriptomic analyses for the genes in the region identified that *ANKRD35* transcription is significantly associated with the chromosome population substructure. Ideogram for the analyzed region showing the transcripts and the four segmental duplication blocks originating the common and minimal deletions associated to the TAR syndrome. The tracks also show the recomb proportion for each population as given by *recombClust* for the 1000 Genomes and for GTEx, demonstrating high reproducibility for the recombination patterns. The recombination differences of the recombination patterns between chromosome subpopulations is also shown. In the lower track, LD between the SNPs in the region and the chromosome subpopulations. SNP eQTLs for *ANKRD35* are depicted in blue and the risk factor for hypertension (rs72704264) is shown in green.

We further asked whether the recombination histories could have a functional role. We tested, using RNA-sequencing data in blood from the GTEx project, if the expression levels of the genes in 1q21.1 were associated with the two different recombination histories. We found a significant differential expression of *ANKRD35* (log fold change = 0.18, *P* = 6.7×10^−4^) and noted that the SNP rs10910843, an eQLTs of *ANKRD35* in blood [32], was in high linkage with the chromosome subpopulations. We additionally found that the SNP rs72704264, a risk factors for hypertension [33], was also in high linkage with the subpopulations, showing likely functional links associated to the different recombination histories.

## Discussion

*recombClust* is the first method to classify chromosomes into different subpopulations based on the inference of the recombination histories along genomic regions. Linkage methods for detecting historic recombination patterns have been important to characterize the distribution of recombination hot-spots between species and ancestries [34–36]. While current methods aim is to robustly estimate the recombination rate between markers by coalescent modeling, accounting for selection and demographic effects, they do not detect recombination variation between individuals. *recombClust* fills this gap, further allowing to test the association between differences in recombination histories with phenotypes.

*recombClust* assumes that there is an inverse relationship between recombination and linkage between genetic markers (SNP-blocks). However, the similarity of the recombination patterns obtained with *recombClust* with those obtained with *FastEPRR* shows that this assumption is not inaccurate. This is because *recombClust* is also the first method to incorporate the spatial correlation of the recombination signal along a genomic region, which other linkage methods do not. Consequently, demographic and selection signals, which induce spatial correlation, are directly extracted from the data (Figures 5–6). Additional analyses are, however, required to identify the nature of different recombination histories and to determine whether they are due to ancestry, selection or the presence of chromosomal rearrangements affecting the recombination within the region. In particular, the method successfully split the groups of chromosomes being selected in the *LCT* locus from those which are not, accurately giving a flat recombination pattern to the group under selection. This is an added advantage with respect to methods like *FastEPRR* in the computation of recombination patterns because *recombClust* explicitly extracts the selection signal from the data by identifying the chromosomes under selection as those with a flat recombination pattern in the locus. Our analyses showed that at the *LCT* locus, the pattern differences between chromosomes groups where large, further suggesting a novel approach in the detection of selection signals.

We have shown that when recombination modifiers are expected to affect a genomic region, such as inversions, *recombClust* can be reliably used to infer its alleles in large population samples. *recombClust* can, for instance, be added to other methods that genotype inversion from SNP data, offering an additional signal derived from recombination patterns [37]. However, we expect that the limitations of these methods also apply to *recombClust*, such as inversions being ancient and not recurrent. Recombination modifiers acting on small targeted sequences that are not expected to show a spatial-extended historic pattern require further methodological developments, like merging the mixture model with coalescent modeling. In general, recombination modifiers whose effects cannot be observed in historical recombination patterns are beyond linkage methods.

We also showed that *recombClust* can detect differences in recombination histories in complex regions prone to non-allelic homologous recombination (NAHR) and, therefore, likely subjected to tight regulation of recombination [30]. We discovered and validated the existence or two recombination histories in the 1q21.1 locus at risk of deleterious syndromes. Detailed analyses are needed to disentangle the nature of the recombination modifiers acting on the region, which can be, for instance, a mixture of genomic rearrangements, epigenetic marks or functional mechanisms regulating double strand breaks that avoid NAHR [30]. In addition, the question arises of whether the recombination between the chromosome subpopulations confers specific risks to deletions and duplications in the offspring. As for the subpopulations’ relation with more common phenotypes, we observed a strong linkage with a risk factor for hypertension showing probable implications of recombination variation with this trait within 1q21.1. We, therefore, showed an approach to measure the impact of different recombination histories on phenotypes, opening a way to study how recombination variation influences traits.

## Methods

### *recombClust* description

We proposed a method to classify chromosomes according to the combinations of SNP alleles across a genomic region that are allowed by different recombination patterns. Consider a situation where two recombination patterns are latent in the chromosome population generating two chromosome subpopulations in a given genomic region (Sup Figure 8). A first subpopulation of chromosomes comprises those that have recombined at any of three given points within the region, and a second subpopulation comprises those that have recombined at any of two other points. In this case, we can see, for instance, that while two specific haplotypes G1 and H1 are compatible with the recombination pattern 1, they are maximally different in mutation content at each SNP variant. In addition, H1 is more similar in mutation content to H2 than G1 is to H1, despite H1 and H2 belonging to different recombination subpopulations. In this work, we proposed the method *recombClust* that first classifies chromosomes into those that have recombined in point between two markers and those that have not, and second, it computes a consensus classification of chromosomes across all points, separating the population of chromosomes according to different recombination patterns along the segment.

#### Mixture model to classify a fraction of recombining chromosomes

The first step of recombClust is the classification of chromosomes that have recombined at one point flanked by two SNP blocks. We therefore propose to model the likelihood that a chromosome in the sample is drawn from a mixture of chromosomes that highly recombined at the point (recomb) and that remained in complete LD (linkage) (Figure 1A). The likelihood is therefore given by a mixture of two latent chromosome groups (recomb/linkage). In the first group, we model the recombination at a point that lies in the sequence interval between a pair of SNP blocks (i=1, 2) of length *L*. Phased SNP alleles are encoded by 0 or 1, the haplotype of a chromosome at block i is a random variable denoted *X_i_* ∈ {0,1}^*L*^ and the haplotype of the joint blocks is the random variable given by the concatenation of the block variables *X*_12_ = *X*_1_ ∘ *X*_2_. Under this model, the recombination completely breaks the LD between the SNP blocks (r^2^ = 0) in the recomb subpopulation and therefore *X*_1_ and *X*_2_ are statistically independent. Therefore, the probability that a chromosome is observed with haplotype *x*_12_ in a chromosome group under recombination is:

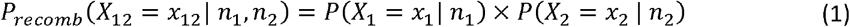

given the haplotype frequencies *n*_1_ and *n*_2_.

For the second chromosome group, we consider that there is no recombination and we model the SNP blocks to be in complete LD (r^2^ = 1). For the chromosomes in the *linkage* group, *X*_1_ and *X*_2_ are completely linked. *X*_2_ can be unambiguously mapped to *X*_1_ (*f* : *X*_2_ → *X*_1_). Under this model, the probability of observing haplotype *x*_12_ is:

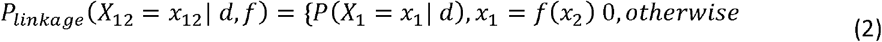

where *d* are the frequencies of *X*_1_.

We define the mixture model with two components, following equations (1) and (2). The model represents a chromosome population with a mixture of *recomb* and *linkage* groups with proportion *π*. We therefore assume that the probability of observing a chromosome with haplotype *x*_12_ is

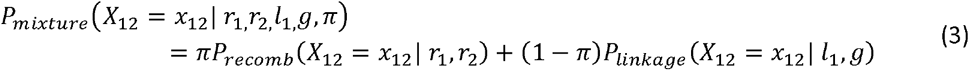

where *r*_1_ and *r*_2_ are the frequencies of haplotypes *X*_1_ and *X*_2_ in *recomb*, *l*_1_ is the haplotype frequencies of *X*_1_ in *linkage*, where *g* is the function linking *X*_2_ to *X*_1_.

Given a set of *m* independent chromosomes (*k* = 1, … *m*), we denote the random variable for the joint blocks over all chromosomes as 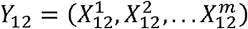 and therefore the likelihoods of observing the data *y*_12_ under the *mixture* model is:

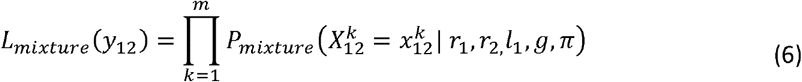

The *mixture* model parameters are determined using an Expectation-Maximization (EM) algorithm. For each chromosome, we define a hidden variable *z_k_∈*{0,1}. This variable indicates if the chromosome belongs to the *recomb* or the *linkage* groups. The EM algorithm updates the model parameters iteratively maximizing the expectation of the data. Given the parameters of the model *ω*, *ω* = (*r*_1_ *r*_2_, *l*_1_, *g*, *π*), we define the probability that chromosome *k* belongs to the *linkage* group, 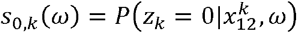. Similarly, the probability that individual *k* belongs to the *recomb* group given *ω* is 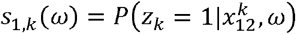. For each *k* the probability of belonging to any group is 1 and, therefore, *s*_0,*k*_(*ω*) + *s*_1,*k*_(*ω*) = 1. In each step of the EM algorithm, we find the value of *ω*’ that maximizes:

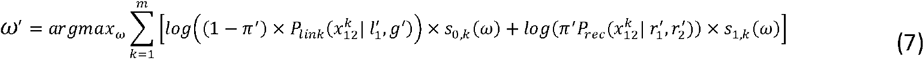

We therefore update the mixture likelihood by *ω*’given by:

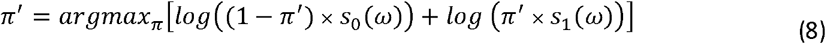

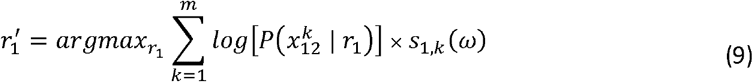

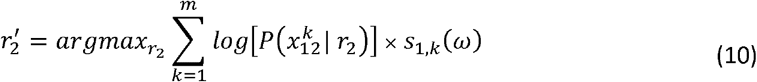

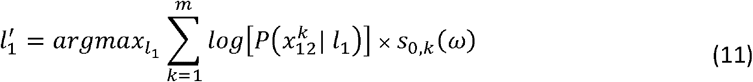

We estimate haplotype frequencies *r*_1_, *r*_2_, and *l*_1_ in close form using Lagrange multipliers, following Sindi *et al*. [38]. In particular, we obtain

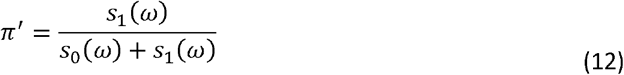

Where *s*_0_(*ω*) and *s*_1_(*ω*) are the probabilities that a chromosome in the population belongs to the *linkage* or the *recomb* groups 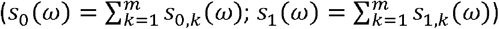. We consider that a chromosome *k* belongs to *recomb* if *s*_1, *k*_ > 0.5. The function *g*’ is defined using a greedy algorithm which sequentially pairs each observed *r*_2_, in decreasing order by their frequency, with the *x*_1_ for which the observed frequency of *x*_12_, is maximum and has not been previously paired. The final *ω*’ is such that its square root difference with the previous estimate is lower than machine precision. In addition, for numerical stability we set the zero in equation 2 to 10^−5^.

#### Clustering of chromosomes into different recombination patterns

Differences in recombination patterns are given by the recombination points in which only a fraction of chromosomes showed historical recombination. In the second step of *recombClust*, a consensus clustering is performed on all the recombination points tested over a genomic region to determine whether individual chromosomes are consistently classified into different recombination patterns. Therefore, to detect a subpopulation of chromosomes across the region based on their recombination patterns, *recombClust* first extensively fits the mixture model between numerous non-overlapping 2-SNP blocks. For each model, the method computes the probability that the chromosomes belong to the *recomb* group. Finally, *recombClust* produces a consensus classification of the chromosomes by clustering the first principal component of the *recomb* probabilities matrix across all mixture models fitted in the genomic region (Figure 1B).

#### Extraction of recombination patterns along a genomic region

We defined *recombClust* recombination patterns as the proportion of chromosomes that have recombined in each subpopulation at different points inside a target region. We started by dividing the target region in non-overlapping windows. In each window, we selected those models overlapping the window. In each model, we assigned a chromosome to *recomb* group if its probability of belonging to the *recomb* group was higher than 0.5. Then, we consider that the chromosome belonged to recombining group in a given window when it was assigned to the *recomb* group in more than half of the models. We defined non-overlapping windows of 50Kb for human ^~^4Mb inversion 8p23.1 and the ^~^1Mb LCT region and of size 20Kb in the 0.4Mb 1q21.1 region.

### Simulation to assess mixture model performance

We evaluated the accuracy of the mixture model to classify individual chromosomes with extensive simulations. 200 instances of a reference scenario were generated and compared with the 200 instances of multiple scenarios under different SNP block and between chromosome group variabilities. For one instance of the reference scenario, we simulated 1000 chromosomes in the *recomb* and the *linkage* groups each, given be the random and full linkage association between a pair of two-SNP blocks, respectively. For the *recomb* group, the chromosome alleles at each SNP were drawn from a binomial distribution whose frequency was independently sampled from a uniform distribution (unif(0.55, 0.95)), assuming no LD within the blocks and between blocks. For the *linkage* group, SNPs within the blocks were independent but the pair of blocks, flanking the recombination point, was in maximum LD. We then considered that the most frequent haplotype for the joint SNP blocks was the same in both subpopulations and given by the SNP alleles with maximum frequency, so the overall linkage in the total population was of D’=1. Different scenarios were obtained by changing the parameters of these simulations, where assessed the performance of the mixture model, given by the accuracy to correctly classify chromosomes into the *recomb/linkage* groups. We first assessed the extent to which the accuracy of the model was affected by the genetic variability between populations, by considering that the differences between the most frequent block-pair haplotypes in each chromosome group was increasingly higher. We did this by changing the number of SNP alleles that were different between the most frequent haplotypes in each group.

We also assessed the influence of within block variability on model accuracy, by taking blocks where the linkage between the SNPs in the block was maximum. This scenario reduces to having blocks of 1 SNP. Finally, we evaluated how the proportion between *recomb* and *linkage* populations affected the mixture model performance. We simulated different scenarios where the proportion of the *recomb* population ranged between 0.1 and 0.9. We test the model the reference scenario and using different initializations for the mixture frequency.

### Performance of *recombClust* to detect chromosomes with different recombination histories

We also evaluated the performance of classifying the chromosomes under different recombination patterns using simulated inversions. As inversion polymorphisms produce chromosomal subpopulations that differ in their recombination patterns, we tested the ability of *recombClust* to detect inversion status in simulated inversions. We simulated an inversion of 800 Kb and a frequency of 20% using *invertFREGENE* [19] to evaluate the mixture model at different recombination points. We varied the inversion length (from 50 Kb to 1 Mb) and inversion frequency (from 0.1 to 0.9) to evaluate the overall *recombClust* performance to call the inversion status of the chromosomes. Each combination of frequency and length was run 100 times. In all simulations, we used the default values of *invertFREGENE* parameters (recombination: 1.25 × 10^−7^, mutation rate: 2.3 × 10^−7^).

### *Drosophila Melanogaster* and human inversions

We tested whether *recombClust* could characterize chromosomal inversions using differences in recombination patterns in *Drosophila Melanogaster* and in humans. We used *recombClust* to infer inversion status of chromosomes for three well known inversions: In(2L)t (2?2225744-13154180, dm6), In(2R)NS (2R:11278659-16163839, dm6) and In(3R)Mo (3R:17232639-24857019). We used SNP data from DGRP2 lines [20, 21], excluding individuals with call rate < 95% and SNPs having any missing or a MAF < 5%, classified the lines into the underlying recombination patterns computed by *recombClust* and compared the classification with experimental inversion genotypes [21].

We used *recombClust* to classify phased chromosomes into underlying recombination patterns within human inversions at 8p23.1 (chr8:8055789-11980649, hg19) and 17q21.31 (chr17:43661775-44372665, hg19). We used SNP phased data from the 1000 Genomes project [25]. We inferred the recombination modifier variants with *recombClust* and compared them with the experimental inversion genotypes available in the *invFEST* repository [22].

### Recombination substructure in the susceptibility region of TAR syndrome

We ran *recombClust* across the region chr1:145.35-145.75Mb characterized by four blocks of segmental duplications. This region is prone to deleterious rearrangements by non-allelic homologous recombination (NAHR), which can lead to the thrombocytopenia-absent radius (TAR) syndrome. We analyzed the 503 European individuals from the 1000 Genomes project and the 528 European individuals of Genotype-Tissue Expression (GTEx) project [39]. We obtained GTEx data from dbGAP (accession code: phs000424.v7.p2), we phased it with *SHAPEIT* [40] and we selected those individuals classified as European by *peddy* [41] with a probability higher than 0.9. In the *recombClust* analysis, we included SNPs with a MAF > 0.05 and performed the consensus clustering across the detected points with a hierarchical clustering. In GTEx, we used the first two PCs of the chromosome subpopulation probabilities while in 1000 Genomes we only used the second PC. We tested Hardy-Weinberg equilibrium using *SNPassoc* [42].

We studied whether the chromosome genotypes, derived from the chromosome subpopulations, were associated with gene expression and phenotype differences between individuals. We evaluated the association with gene expression in whole blood using GTEx data, using the gene raw counts from *recount2* (33). For each tissue, we removed genes with less than 10 counts in more than 90% of the samples. We tested the association between the chromosome alleles and gene expression, applying a robust linear regression with *limma* [43] to log_2_CPM values obtained with *voom* [44]. We included sex, platform, top three genome-wide principal components and variables from PEER as covariates.

## Supplementary material

Additional File 1: .pdf. Supplementary Figures and Tables. Sup Figures 1–8, Sup Table 1

## Acknowledgments

The authors would like to express their gratitude to the Supercomputing and Bioinnovation Center (SCBI) of the University of Malaga (Spain) for their support and resources. The Genotype-Tissue Expression (GTEx) Project was supported by the Common Fund of the Office of the Director of the National Institutes of Health, and by NCI, NHGRI, NHLBI, NIDA, NIMH, and NINDS. GTEx data were obtained from: the GTEx Portal on 06/07/2018 and dbGaP accession number phs000424.v7.p2 on 12/05/2017.

---

**Sup Figure 1:**
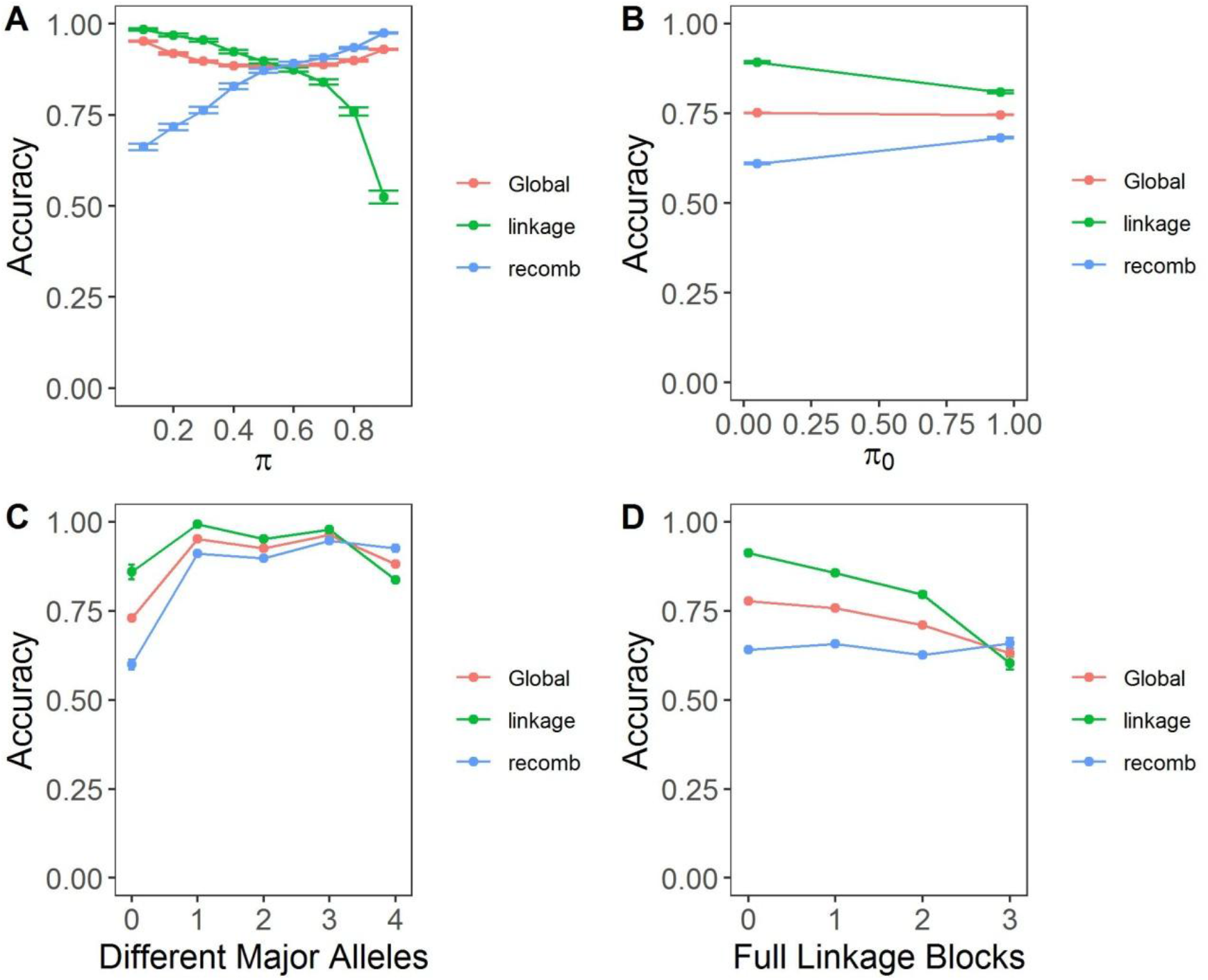
Evaluation of the mixture model under different simulated scenarios. Each plot contains the accuracy for classifying all chromosomes and the accuracies to classify *recomb* or *linkage* chromosomes (i.e. sensibility). π: proportion of chromosomes belonging to *recomb* population. π_0_: *recomb* proportion initial value. Different Major Alleles: SNPs having different major alleles in *recomb* and linkage populations. Full Linkage Blocks: Number of blocks without variability, i.e. they can be considered as a single SNP. Accuracies contain the mean and standard error computed from 200 simulations of each scenario.

**Sup Figure 2:**
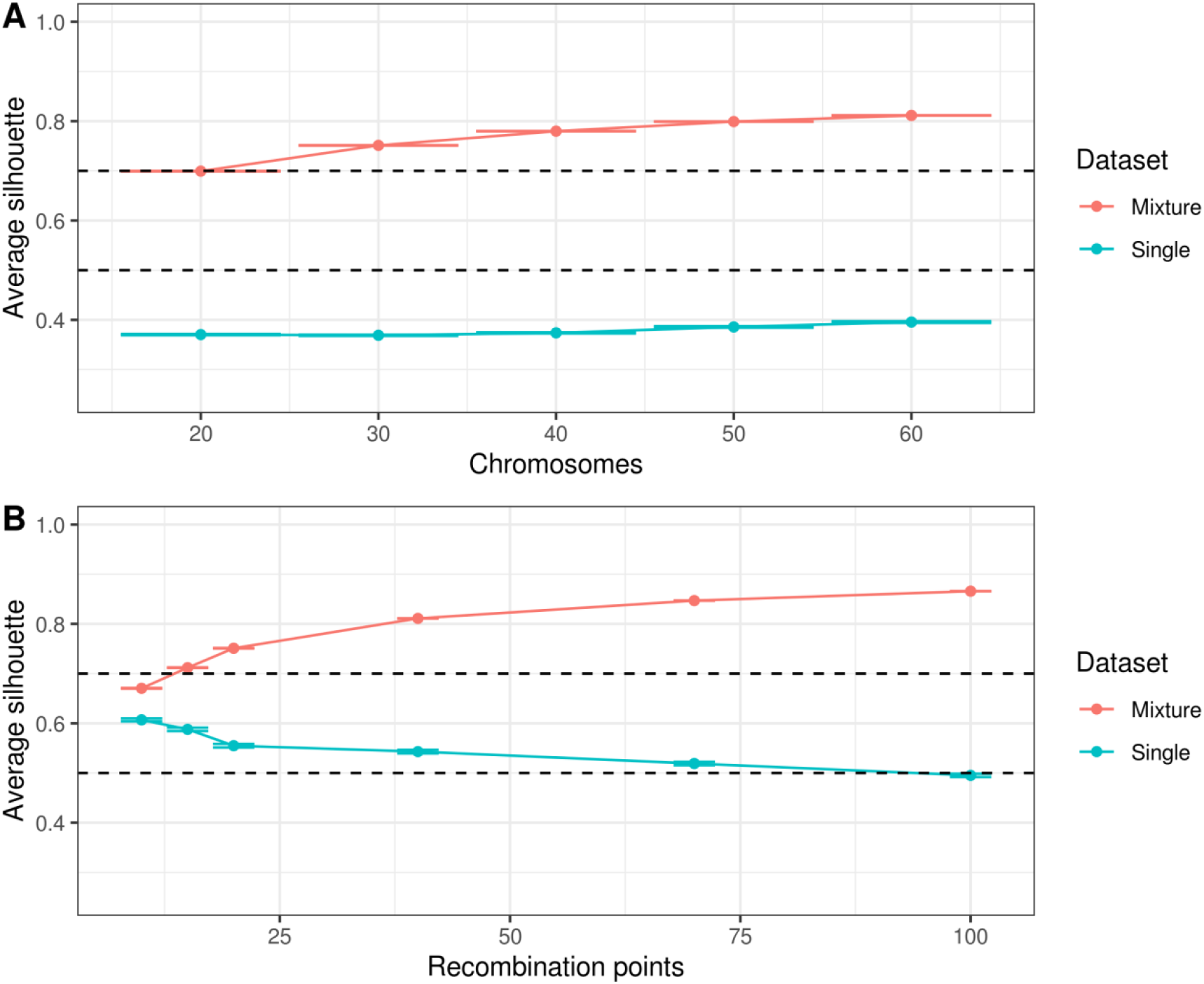
Accuracy of *recombClust* to detect datasets with a population mixture. Average silhouette value indicates how reliable was recombClust clustering in a given dataset. Dashed lines mark critical cut-off (>0.7: very reliable structure; 0.5-0.7: reliable structure; <0.5: unreliable clustering). Mixture datasets contains two subpopulations with non-overlapping recombination points, while single datasets contain one population. Average silhouette values contain the mean and standard error computed from 1000 simulations of each scenario.

**Sup Figure 3:**
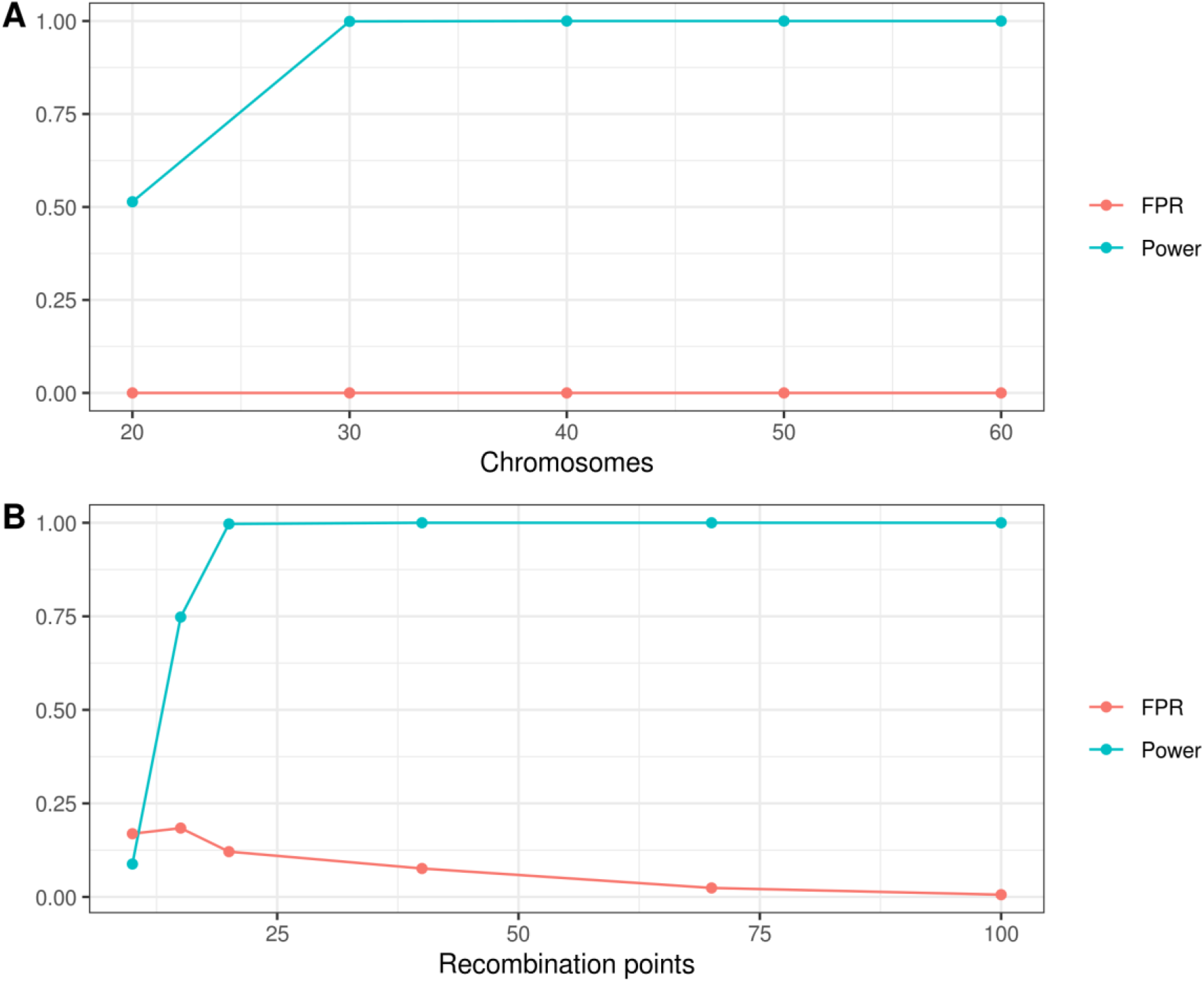
*recombClust* evaluation in simulated datasets with a known population mixture. We computed FPR and power based on simulated datasets. Half of the datasets contained a mixture population and the other half one population. A dataset with an average silhouette value of 0.7 was considered as supporting a mixture population by recombClust. Each point is the FPR and power of 2000 simulations.

**Sup Figure 4:**
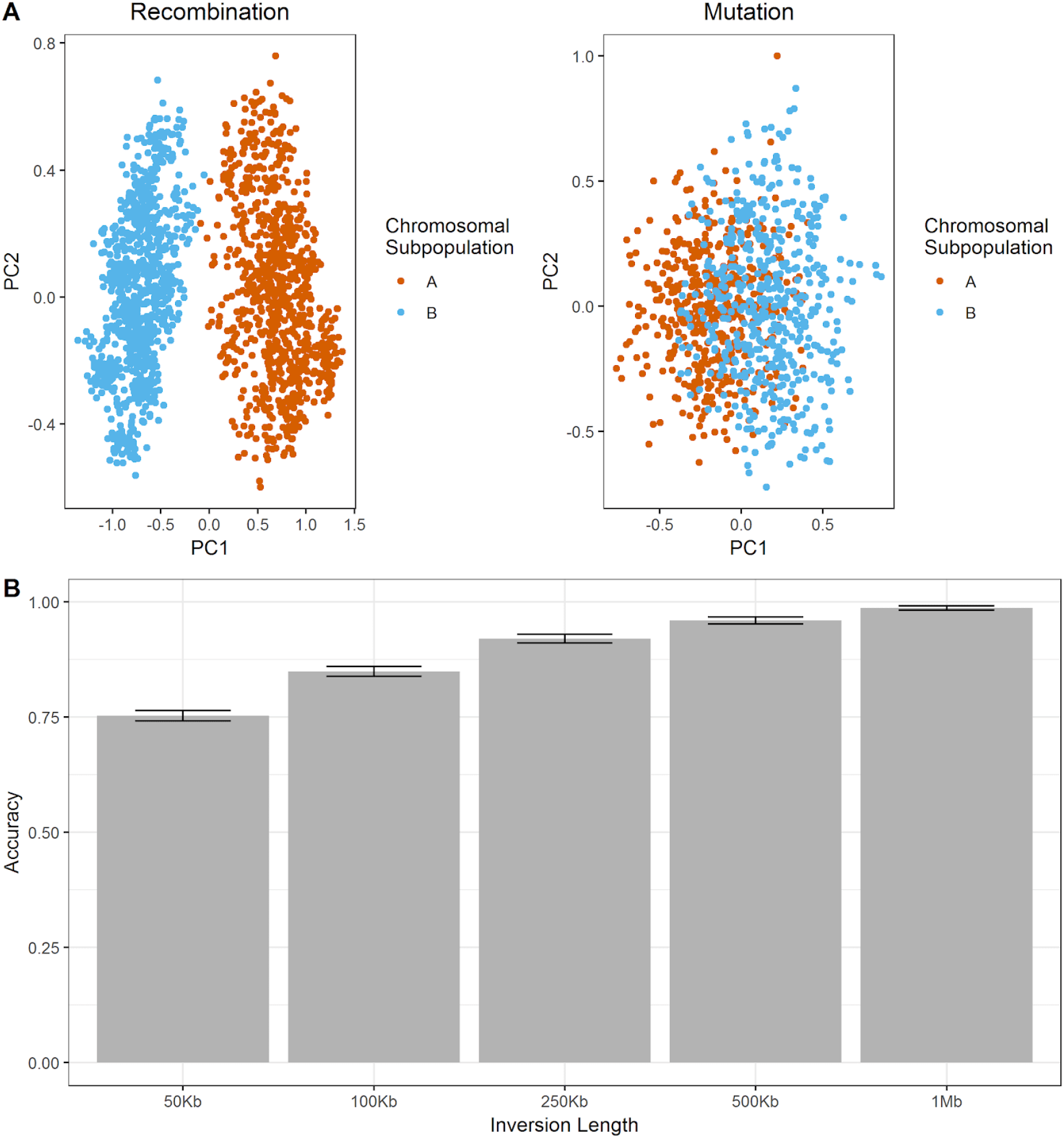
recombClust accuracy for detecting subpopulations with different recombination patterns. A-left) Detection of recombination patterns on simulated data by clusters on the PCs of the prediction matrix across mixture models on 10 recombination points. Five different recombination points were simulated for each subpopulation A and B, where the other subpopulation remained in linkage. The first PC shows a clear separation of the subpopulations. A-right) To test mutation differences between the subpopulations, we computed the PC for the genotype matrix of the markers flanking the 10 recombination points. In this case the PCs did not showed a clear separation between the chromosome subpopulations. B) The figure shows the match between the chromosome subpopulations as obtained by recombClust and inversion status of chromosomes, for 9,000 simulated inversions at a given size (1000 simulations at 9 different inversion frequencies). The figure shows the mean accuracy and standard error. recombClust identifies inversion status by recombination differences with high accuracy, particularly for inversions > 250Kb.

**Sup Figure 5:**
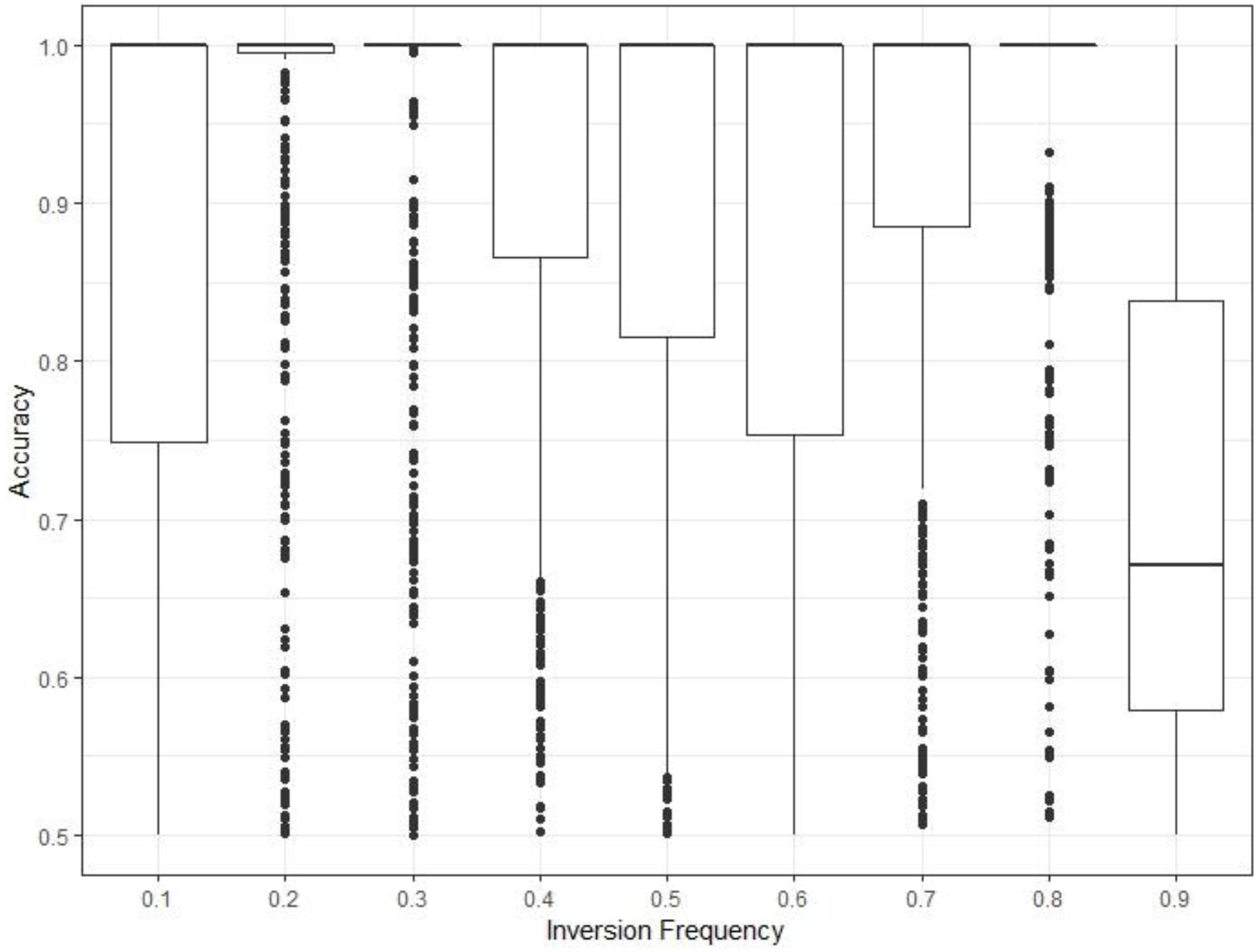
*recombClust* accuracy for different inversion frequencies. Accuracy is the proportion of phased chromosomes correctly classified. Each boxplot includes 500 simulations.

**Sup Figure 6:**
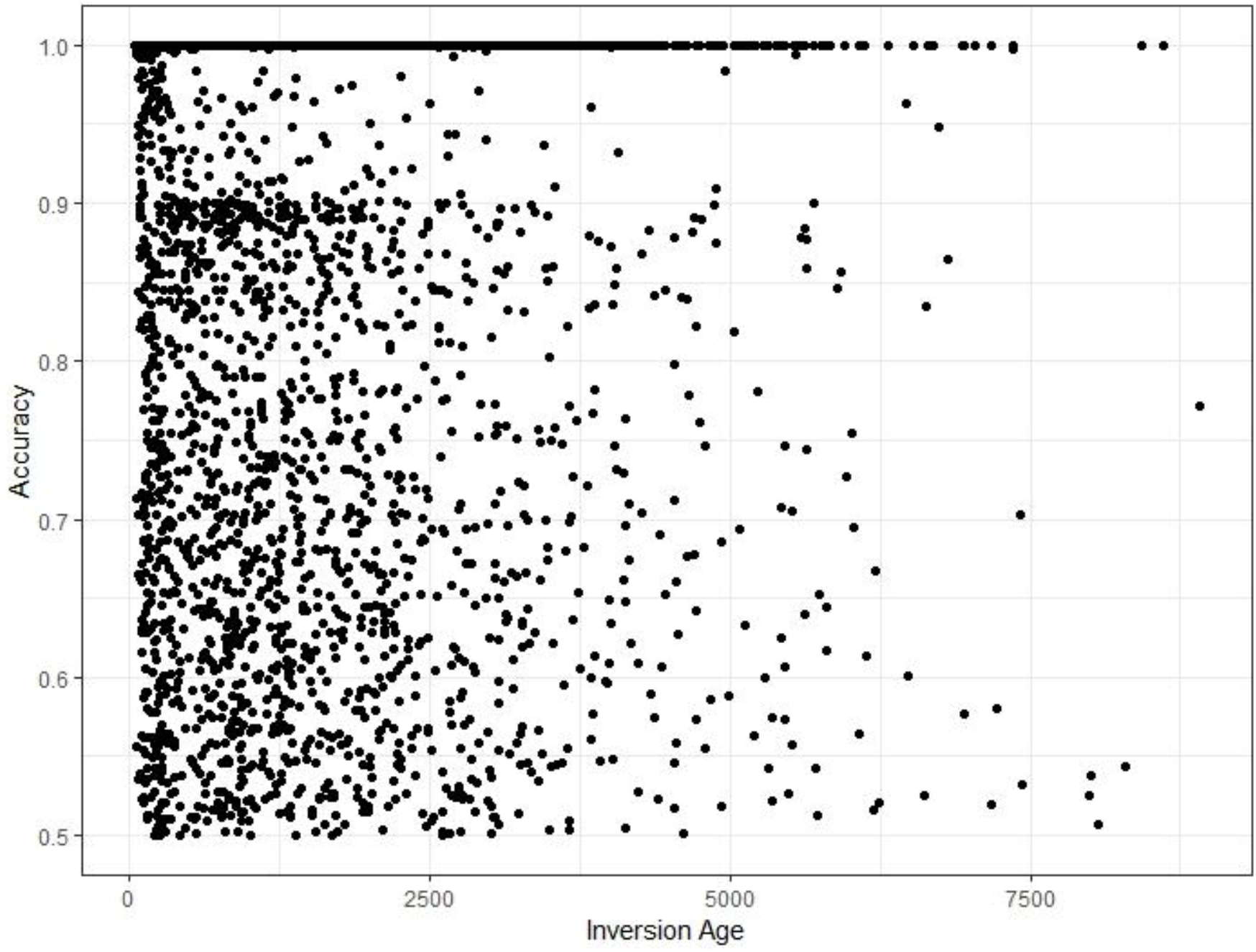
*recombClust* accuracy for different inversion ages. Accuracy is the proportion of phased chromosomes correctly classified. Each point is the accuracy of an independent simulation.

**Sup Figure 7:**
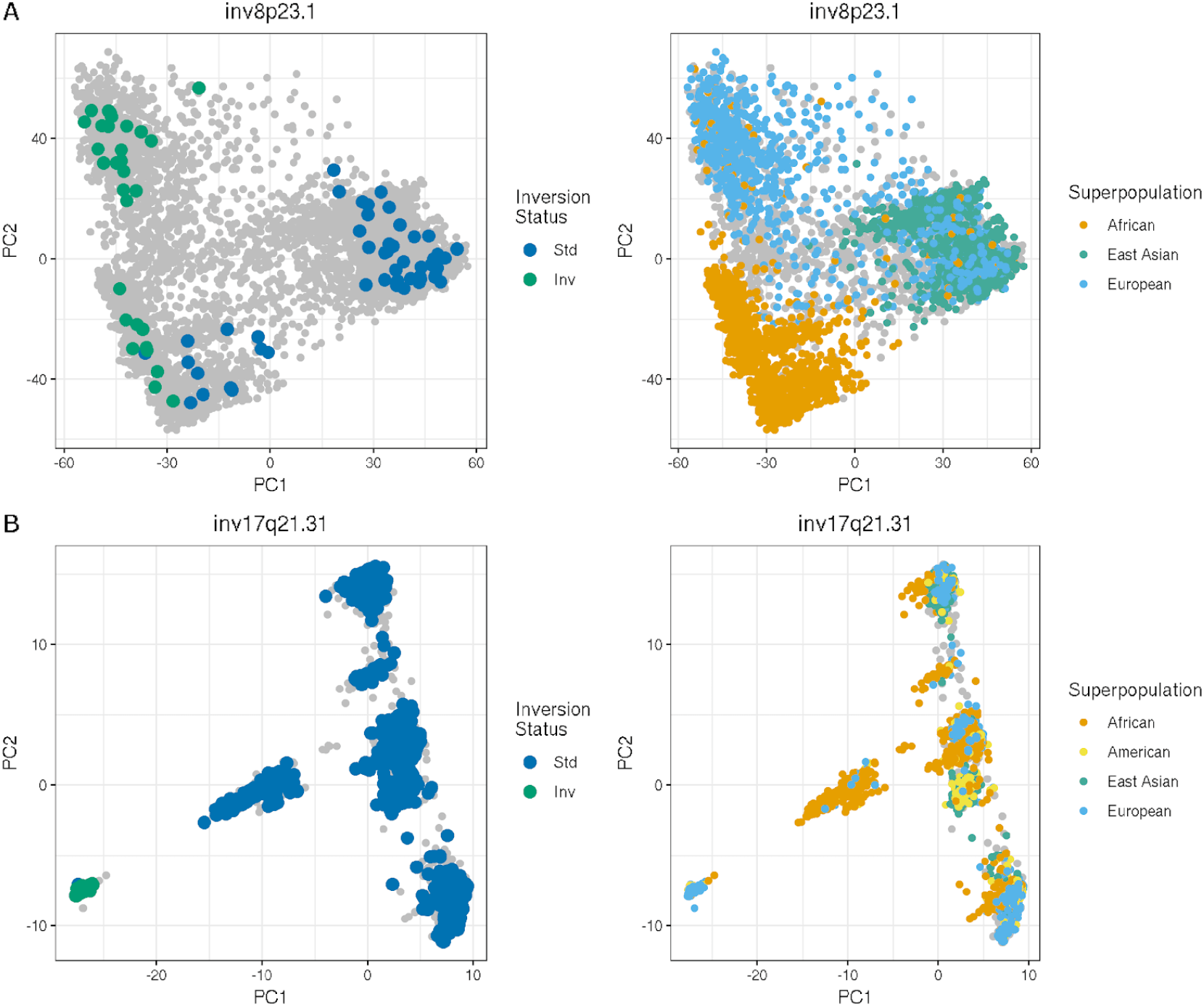
Identification of chromosomal subpopulations of different ancestries from differences in the recombination patterns within two inversions. The figures show the first two PCA components for the all mixture model predictions at numerous recombination points across inv-8p23.1 and inv-17q21.31, computed for all 1000 Genomes ancestries. Chromosomes are clearly separated by inversion status (Std, Inv) and ancestry. For inv-8p23.1 clear ancestral groups are identified within inversion status whereas ancestry is mixed within each inv-17q21.31 status. Colored points indicate experimentally validated observations of inversion status and ancestry.

**Sup Figure 8:**
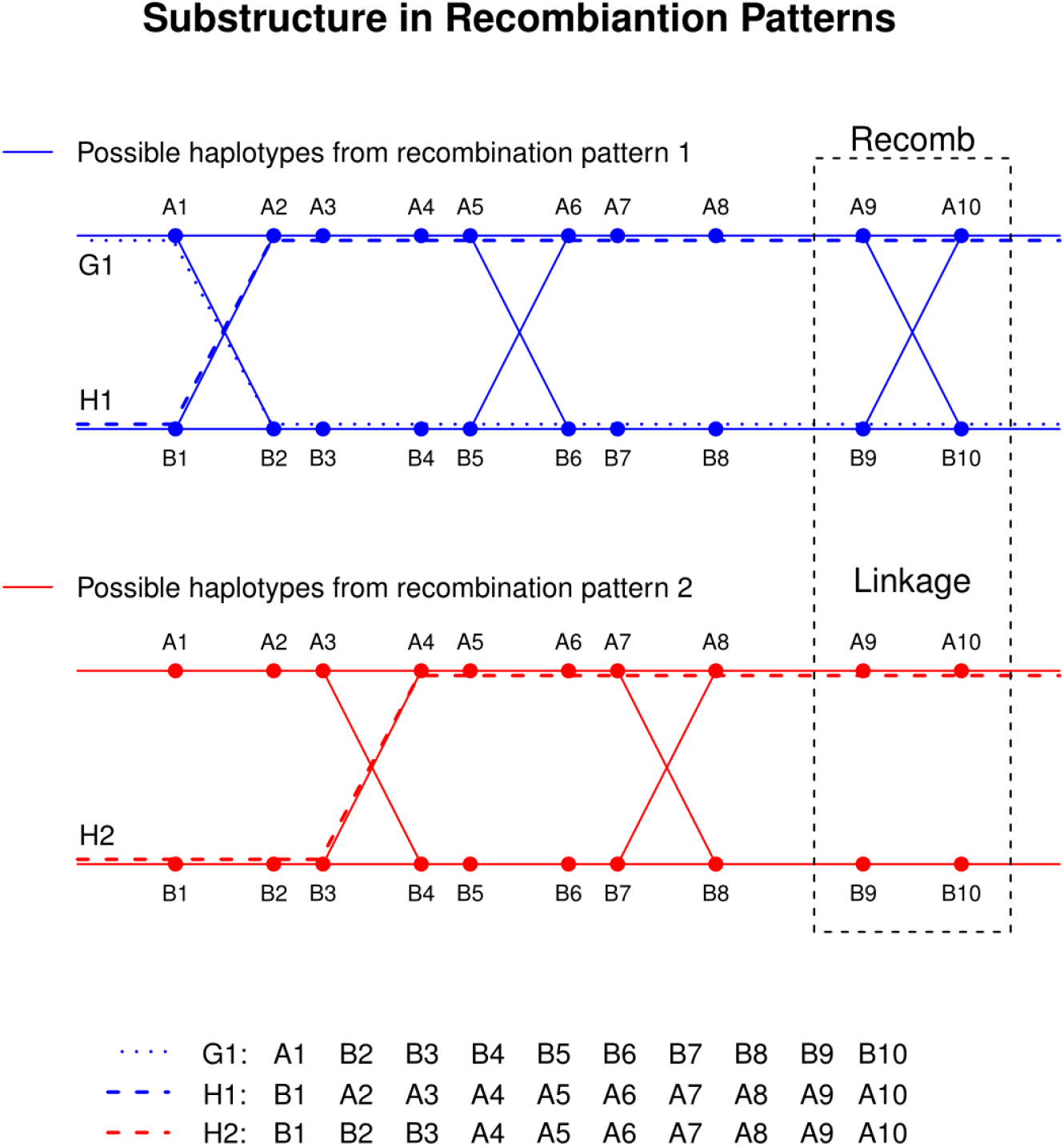
Representation of two chromosomal subpopulations with different recombination patterns in a genomic segment. Lines represent the possible chromosomes present in population 1 (blue) and population 2 (red). Each SNP has two alleles (A and B) and is labelled with a number. Recombination points are placed between SNPs where A and B alleles are joined by a line. G1 and H1 are two possible chromosomes from population 1 and H2 is one of the possible chromosomes from population 2. The dotted box contains a recombination point present in population 1 but not in population 2.

**Sup Table 1:** recombClust allele frequencies in LCT locus for different European populations. Allele 1 is more frequent in all populations by TSI, the only population that do not show a selection mark based on iHS. CEU: Utah residents (CEPH) with Northern and Western European ancestry. FIN: Finnish in Finland. GBR: British in England and Scotland. IBS: Iberian populations in Spain. TSI: Toscani in Italy.

|  | Allele 1 Frequency | Allele 2 Frequency |
| --- | --- | --- |
| CEU | 79.8% | 20.2% |
| FIN | 65.7% | 34.3% |
| GBR | 74.2% | 25.8% |
| IBS | 59.8% | 40.2% |
| TSI | 28.5% | 71.5% |

## References

1. Alves I, Houle AA, Hussin JG, Awadalla P: The impact of recombination on human mutation load and disease. Philos Trans R Soc B Biol Sci 2017, 372:20160465.

2. Kong A, Thorleifsson G, Gudbjartsson DF, Masson G, Sigurdsson A, Jonasdottir A, Walters GB, Jonasdottir A, Gylfason A, Kristinsson KT, Gudjonsson SA, Frigge ML, Helgason A, Thorsteinsdottir U, Stefansson K: Fine-scale recombination rate differences between sexes, populations and individuals. Nature 2010, 467:1099–1103.

3. Alves JM, Chikhi L, Amorim A, Lopes AM: The 8p23 Inversion Polymorphism Determines Local Recombination Heterogeneity across Human Populations. Genome Biol Evol 2014, 6:921–930.

4. McVean GAT, Myers SR, Hunt S, Deloukas P, Bentley DR, Donnelly P: The Fine-Scale Structure of Recombination Rate Variation in the Human Genome. Science (80-) 2004, 304:581–584.

5. Stumpf MPH, McVean GAT: Estimating recombination rates from population-genetic data. Nat Rev Genet 2003, 4:959–68.

6. Auton A, McVean G: Estimating Recombination Rates from Genetic Variation in Humans. In Methods in molecular biology (Clifton, N.J.). Volume 856; 2012:217–237.

7. Nei M: Modification of linkage intensity by natural selection. Genetics 1967, 57:625–41.

8. Feldman MW, Otto SP, Christiansen FB: POPULATION GENETIC PERSPECTIVES ON THE EVOLUTION OF RECOMBINATION. Annu Rev Genet 1996, 30:261–295.

9. Kirkpatrick M, Barton N: Chromosome inversions, local adaptation and speciation. Genetics 2006, 173:419–34.

10. Stefansson H, Helgason A, Thorleifsson G, Steinthorsdottir V, Masson G, Barnard J, Baker A, Jonasdottir A, Ingason A, Gudnadottir VG, Desnica N, Hicks A, Gylfason A, Gudbjartsson DF, Jonsdottir GM, Sainz J, Agnarsson K, Birgisdottir B, Ghosh S, Olafsdottir A, Cazier J-B, Kristjansson K, Frigge ML, Thorgeirsson TE, Gulcher JR, Kong A, Stefansson K: A common inversion under selection in Europeans. Nat Genet 2005, 37:129–137.

11. Puig M, Casillas S, Villatoro S, Cáceres M: Human inversions and their functional consequences. Brief Funct Genomics 2015, 14:369–79.

12. Hussin J, Sinnett D, Casals F, Idaghdour Y, Bruat V, Saillour V, Healy J, Grenier J-C, de Malliard T, Busche S, Spinella J-F, Lariviere M, Gibson G, Andersson A, Holmfeldt L, Ma J, Wei L, Zhang J, Andelfinger G, Downing JR, Mullighan CG, Awadalla P: Rare allelic forms of PRDM9 associated with childhood leukemogenesis. Genome Res 2013, 23:419–430.

13. Thacker D, Keeney S: Homologous Recombination During Meiosis. In DNA Replication, Recombination, and Repair. Tokyo: Springer Japan; 2016:131–151.

14. Myers S, Bottolo L, Freeman C, McVean G, Donnelly P: A Fine-Scale Map of Recombination Rates and Hotspots Across the Human Genome. Science (80-) 2005, 310:321–324.

15. Coop G, Przeworski M: An evolutionary view of human recombination. Nat Rev Genet 2007, 8:23–34.

16. Rosenfeld JA, Traylor RN, Schaefer GB, McPherson EW, Ballif BC, Klopocki E, Mundlos S, Shaffer LG, Aylsworth AS, 1q21.1 Study Group 1q21.1 Study: Proximal microdeletions and microduplications of 1q21.1 contribute to variable abnormal phenotypes. Eur J Hum Genet 2012, 20:754–61.

17. Mefford HC, Sharp AJ, Baker C, Itsara A, Jiang Z, Buysse K, Huang S, Maloney VK, Crolla JA, Baralle D, Collins A, Mercer C, Norga K, de Ravel T, Devriendt K, Bongers EMHF, de Leeuw N, Reardon W, Gimelli S, Bena F, Hennekam RC, Male A, Gaunt L, Clayton-Smith J, Simonic I, Park SM, Mehta SG, Nik-Zainal S, Woods CG, Firth H V., et al.: Recurrent Rearrangements of Chromosome 1q21.1 and Variable Pediatric Phenotypes. N Engl J Med 2008, 359:1685–1699.

18. Kaufman L, Rousseeuw PJ: Finding Groups in Data⍰: An Introduction to Cluster Analysis. Wiley-Interscience; 1990.

19. O’Reilly PF, Coin LJM, Hoggart CJ: invertFREGENE: software for simulating inversions in population genetic data. Bioinformatics 2010, 26:838–840.

20. Mackay TFC, Richards S, Stone EA, Barbadilla A, Ayroles JF, Zhu D, Casillas S, Han Y, Magwire MM, Cridland JM, Richardson MF, Anholt RRH, Barrón M, Bess C, Blankenburg KP, Carbone MA, Castellano D, Chaboub L, Duncan L, Harris Z, Javaid M, Jayaseelan JC, Jhangiani SN, Jordan KW, Lara F, Lawrence F, Lee SL, Librado P, Linheiro RS, Lyman RF, et al.: The Drosophila melanogaster Genetic Reference Panel. Nature 2012, 482:173–178.

21. Huang W, Massouras A, Inoue Y, Peiffer J, Ràmia M, Tarone AM, Turlapati L, Zichner T, Zhu D, Lyman RF, Magwire MM, Blankenburg K, Carbone MA, Chang K, Ellis LL, Fernandez S, Han Y, Highnam G, Hjelmen CE, Jack JR, Javaid M, Jayaseelan J, Kalra D, Lee S, Lewis L, Munidasa M, Ongeri F, Patel S, Perales L, Perez A, et al.: Natural variation in genome architecture among 205 Drosophila melanogaster Genetic Reference Panel lines. Genome Res 2014, 24:1193–1208.

22. Martínez-Fundichely A, Casillas S, Egea R, Ràmia M, Barbadilla A, Pantano L, Puig M, Cáceres M: InvFEST, a database integrating information of polymorphic inversions in the human genome. Nucleic Acids Res 2014, 42:D1027–D1032.

23. Gao F, Ming C, Hu W, Li H: New Software for the Fast Estimation of Population Recombination Rates (FastEPRR) in the Genomic Era. G3 (Bethesda) 2016, 6:1563–1571.

24. Jeffreys AJ, Neumann R: The rise and fall of a human recombination hot spot. Nat Genet 2009, 41:625–9.

25. Sudmant PH, Rausch T, Gardner EJ, Handsaker RE, Abyzov A, Huddleston J, Zhang Y, Ye K, Jun G, Fritz MH-Y, Konkel MK, Malhotra A, Stütz AM, Shi X, Casale FP, Chen J, Hormozdiari F, Dayama G, Chen K, Malig M, Chaisson MJP, Walter K, Meiers S, Kashin S, Garrison E, Auton A, Lam HYK, Mu XJ, Alkan C, Antaki D, et al.: An integrated map of structural variation in 2,504 human genomes. Nature 2015, 526:75–81.

26. Steinberg KM, Antonacci F, Sudmant PH, Kidd JM, Campbell CD, Vives L, Malig M, Scheinfeldt L, Beggs W, Ibrahim M, Lema G, Nyambo TB, Omar SA, Bodo J-M, Froment A, Donnelly MP, Kidd KK, Tishkoff SA, Eichler EE: Structural diversity and African origin of the 17q21.31 inversion polymorphism. Nat Genet 2012, 44:872–80.

27. Murga-Moreno J, Coronado-Zamora M, Bodelón A, Barbadilla A, Casillas S: PopHumanScan: the online catalog of human genome adaptation. Nucleic Acids Res 2019, 47:D1080–D1089.

28. Bhérer C, Campbell CL, Auton A: Refined genetic maps reveal sexual dimorphism in human meiotic recombination at multiple scales. Nat Commun 2017, 8:14994.

29. Albers CA, Paul DS, Schulze H, Freson K, Stephens JC, Smethurst PA, Jolley JD, Cvejic A, Kostadima M, Bertone P, Breuning MH, Debili N, Deloukas P, Favier R, Fiedler J, Hobbs CM, Huang N, Hurles ME, Kiddle G, Krapels I, Nurden P, Ruivenkamp CAL, Sambrook JG, Smith K, Stemple DL, Strauss G, Thys C, van Geet C, Newbury-Ecob R, Ouwehand WH, et al.: Compound inheritance of a low-frequency regulatory SNP and a rare null mutation in exon-junction complex subunit RBM8A causes TAR syndrome. Nat Genet 2012, 44:435–439.

30. Sasaki M, Lange J, Keeney S: Genome destabilization by homologous recombination in the germ line. Nat Rev Mol Cell Biol 2010, 11:182–195.

31. Klopocki E, Schulze H, Strauß G, Ott C-E, Hall J, Trotier F, Fleischhauer S, Greenhalgh L, Newbury-Ecob RA, Neumann LM, Habenicht R, König R, Seemanova E, Megarbane A, Ropers H-H, Ullmann R, Horn D, Mundlos S: Complex Inheritance Pattern Resembling Autosomal Recessive Inheritance Involving a Microdeletion in Thrombocytopenia–Absent Radius Syndrome. Am J Hum Genet 2007, 80:232–240.

32. Westra H-J, Peters MJ, Esko T, Yaghootkar H, Schurmann C, Kettunen J, Christiansen MW, Fairfax BP, Schramm K, Powell JE, Zhernakova A, Zhernakova D V, Veldink JH, Van den Berg LH, Karjalainen J, Withoff S, Uitterlinden AG, Hofman A, Rivadeneira F, ’t Hoen PAC, Reinmaa E, Fischer K, Nelis M, Milani L, Melzer D, Ferrucci L, Singleton AB, Hernandez DG, Nalls MA, Homuth G, et al.: Systematic identification of trans eQTLs as putative drivers of known disease associations. Nat Genet 2013, 45:1238–1243.

33. Evangelou E, Warren HR, Mosen-Ansorena D, Mifsud B, Pazoki R, Gao H, Ntritsos G, Dimou N, Cabrera CP, Karaman I, Ng FL, Evangelou M, Witkowska K, Tzanis E, Hellwege JN, Giri A, Velez Edwards DR, Sun Y V., Cho K, Gaziano JM, Wilson PWF, Tsao PS, Kovesdy CP, Esko T, Mägi R, Milani L, Almgren P, Boutin T, Debette S, Ding J, et al.: Genetic analysis of over 1 million people identifies 535 new loci associated with blood pressure traits. Nat Genet 2018, 50:1412–1425.

34. Laayouni H, Montanucci L, Sikora M, Melé M, Dall’Olio GM, Lorente-Galdos B, McGee KM, Graffelman J, Awadalla P, Bosch E, Comas D, Navarro A, Calafell F, Casals F, Bertranpetit J: Similarity in Recombination Rate Estimates Highly Correlates with Genetic Differentiation in Humans. PLoS One 2011, 6:e17913.

35. Smukowski CS, Noor MAF: Recombination rate variation in closely related species. Heredity (Edinb) 2011,107:496–508.

36. Winckler W, Myers SR, Richter DJ, Onofrio RC, McDonald GJ, Bontrop RE, McVean GAT, Gabriel SB, Reich D, Donnelly P, Altshuler D: Comparison of Fine-Scale Recombination Rates in Humans and Chimpanzees. Science (80-) 2005, 308:107–111.

37. Cáceres A, González JR: Following the footprints of polymorphic inversions on SNP data: from detection to association tests. Nucleic Acids Res 2015:1–11.

38. Sindi SS, Raphael BJ: Identification and Frequency Estimation of Inversion. J Comput Biol 2010, 17:517–531.

39. Lonsdale J, Thomas J, Salvatore M, Phillips R, Lo E, Shad S, Hasz R, Walters G, Garcia F, Young N, Foster B, Moser M, Karasik E, Gillard B, Ramsey K, Sullivan S, Bridge J, Magazine H, Syron J, Fleming J, Siminoff L, Traino H, Mosavel M, Barker L, Jewell S, Rohrer D, Maxim D, Filkins D, Harbach P, Cortadillo E, et al.: The Genotype-Tissue Expression (GTEx) project. Nat Genet 2013, 45:580–585.

40. Delaneau O, Zagury J-F, Marchini J: Improved whole-chromosome phasing for disease and population genetic studies. Nat Methods 2013, 10:5–6.

41. Pedersen BS, Quinlan AR: Who’s Who? Detecting and Resolving Sample Anomalies in Human DNA Sequencing Studies with Peddy. Am J Hum Genet 2017, 100:406–413.

42. Gonzalez JR, Armengol L, Sole X, Guino E, Mercader JM, Estivill X, Moreno V: SNPassoc: an R package to perform whole genome association studies. Bioinformatics 2007, 23:654–655.

43. Ritchie ME, Phipson B, Wu D, Hu Y, Law CW, Shi W, Smyth GK: limma powers differential expression analyses for RNA-sequencing and microarray studies. Nucleic Acids Res 2015, 43:e47.

44. Law CW, Chen Y, Shi W, Smyth GK: voom: Precision weights unlock linear model analysis tools for RNA-seq read counts. Genome Biol 2014, 15:R29.

